# Magnetic compass orientation behaviour of Eurasian blackcaps at the predicted 110–120 MHz upper cut-off frequency for radiofrequency-field effects

**DOI:** 10.64898/2026.05.21.726905

**Authors:** Bo Leberecht, Baladev Satish, Leonard Schwigon, Lalitha Venkatraman, Lisa Borowsky, Julia Orthmann, Florian Ippen, Joe Wynn, P. J. Hore, Henrik Mouritsen

## Abstract

Night-migratory songbirds use the Earth’s magnetic field to guide their migratory journeys. Most evidence suggests that the magnetic compass sensor is a flavin-tryptophan radical pair whose operation is disrupted by broadband radiofrequency (RF) fields at frequencies up to ∼116 MHz. Here, we test whether broadband 110–120 MHz RF fields affect the magnetic orientation behaviour of night-migratory Eurasian blackcaps (*Sylvia atricapilla*). We found that the birds oriented in their expected migratory direction in the natural geomagnetic field (NMF) control condition and that they turned their orientation ∼120° when the field was rotated 120° horizontally (changed magnetic field, CMF). When they were exposed to broadband 110–120 MHz RF fields, the birds continued to orient in the appropriate direction in the NMF. In the CMF, we found orientation behaviour that was not consistent with the expected direction. We conclude that the birds could still orient when exposed to 110–120 MHz fields, but speculate that their magnetic orientation capabilities might have been somewhat reduced compared to the control condition. We suggest that 110–120 MHz could represent a grey zone of reduced magnetic orientation capability centred at the cut-off frequency predicted for a flavin-based radical pair (probably in the flavoprotein cryptochrome).

## 1 Introduction

Every year, night-migratory songbirds travel from their breeding quarters to their wintering quarters and back. During their migration, one of the cues they use is a magnetic compass based on the inclination of the geomagnetic field [1–6]. The magnetic compass is light-dependent [6–8] and the currently leading hypothesis suggests that it is based on a so-called radical-pair mechanism [6,9–13]. Magnetic compass information is detected in the retina and transmitted via the visual thalamofugal pathway to a small part of the visual wulst in the avian forebrain named Cluster N [14–17]. When Cluster N is lesioned, birds are unable to use magnetic compass information, whereas the function of their other visual compasses, the star compass and the sun compass, is unaffected by the lesions [14,16–18].

Night-migratory songbirds have been reported to be unable to use their magnetic compass in the presence of weak single-frequency radiofrequency (RF) electromagnetic fields [11,19–24]. However, broadband RF fields ranging from 450 kHz to at least 80 MHz seem to have a significantly larger disruptive effect on the avian magnetic compass than single-frequency fields [25–30]. A magnetic particle-based sensor [31] has also been proposed, but should not be affected by RF fields of such low intensity and high frequency and hence cannot explain the observed disruptive RF field effects [11,32]. In contrast, disruptive effects of broadband RF fields are qualitatively consistent with a radical-pair-based mechanism [9,11,33].

The only proteins currently known to form radical pairs upon photo-activation in the avian retina are cryptochromes (Cry) [9,11,13,34–41]. To date, three Cry genes have been documented in the songbird retina [42], each with two known splice variants (a and b): Cry1a [37,39,43,44], Cry1b [43,45,46], Cry2a [37,43,47–49], Cry2b [48,50], Cry4a [13,40,42,51–53], and Cry4b [47,54]. Except for Cry4b [54], all variants were documented both on the mRNA and protein level. Cry4a shows seasonal rhythmicity, i.e. an increased expression during the avian migratory seasons [40,47].

The radical-pair mechanism hypothesis is based on the light-induced formation of radical pairs [9]. In many, but not all, cryptochromes the non-covalently bound co-factor flavin adenine dinucleotide (FAD) absorbs the light and is one of the suggested radical partners [11,13,55]. Therefore, if FAD is not bound, a cryptochrome cannot be magnetoreceptive. Avian Cry1 and Cry2 seemingly bind FAD very poorly or not at all [50,56–58] and are therefore unlikely to be magnetic field sensors. Cry4b is probably never translated to protein *in vivo*, and when expressed *in vitro* does not bind FAD [54]. In contrast, Cry4a of night-migratory songbirds, chickens, and pigeons stoichiometrically binds FAD [13,36,50,53,55,58,59] and purified Cry4a from night-migratory European robins, *Erithacus rubecula* (*Er*Cry4a), and other birds has been shown to be magnetically sensitive through *in vitro* experiments [13,55,60]. In the avian retina, Cry4a interacts with a membrane-associated, cone-specific interaction partner, the α-subunit of a particular G-protein, which could anchor Cry4a to the membrane (needed for directional responses; see also [61]) and/or be its first interaction partner for potential signal transduction [62,63].

The currently supported radical-pair mechanism in *Er*Cry4a starts with photoexcitation of the FAD co-factor, which initiates the transfer of four electrons along a chain of four tryptophans (Trp), forming a radical pair sensitive to a magnetic field [13,64]. Similar electron transport chains have also been found in other cryptochromes from other organisms [36,41,55,60,65–68]. The magnetic sensitivity of the radical pair arises from interactions between the spins of the unpaired electrons, the ambient magnetic field, and internal (hyperfine) interactions with magnetic nuclei (^1^H and ^14^N atoms in the radicals [9]).

Experimentally and theoretically, it is clear that RF electromagnetic fields can affect radical pair reactions [69]. If one of the radicals had no hyperfine interactions with nearby magnetic nuclei, the radical pair should be most sensitive by far to a particular radiofrequency (the so-called Larmor frequency: ∼1.4 MHz in a static Earth-strength magnetic field of ∼50,000 nT). Even a few hyperfine interactions in both radicals would substantially smear out this effect [9,29,70]. Furthermore, the two radicals would have to have a tiny dipolar interaction, which would only be possible if the radicals were more than about 4 nm apart. If they were that far apart, back electron transfer would be negligibly slow and the magnetic field effect therefore vanishingly small. Thus, specific effects exclusively at the Larmor frequency are extremely unlikely [9]. In general, the greater the number of hyperfine interactions, the wider is the range of radiofrequencies that can potentially disrupt the radical pair mechanism [9,27,29,70].

Some studies have suggested that the avian radical pair could comprise a FAD radical and a counter-radical that had few or no hyperfine interactions [19,20,25,71,72] which, in theory, could be more sensitive to the geomagnetic field than a sensor based on a FAD–Trp radical pair [9,73]. It has been suggested that superoxide (O_2_^•−^) could theoretically act as a counter-radical devoid of hyperfine interactions, but this is highly unlikely due to the extremely short spin-coherence lifetime of O_2_^•−^-containing radical pairs ([70,74,75]; see also [76]). Furthermore, it is almost impossible to imagine a radical with no hyperfine interactions in a cell [6], because the vast majority of organic radicals contain several ^1^H and/or ^14^N atoms, and even if molecules like O_2_^•−^ are considered, there will be hyperfine interactions with ^1^H atoms in surrounding molecules, e.g. water.

The extreme improbability of a dominant Larmor-frequency resonance is consistent with recent studies in which broadband RF fields were shown to have a more disruptive effect on the birds’ magnetic compass than single-frequency fields [26]. Furthermore, several frequency ranges that did not contain the Larmor frequency were found to disrupt magnetic compass orientation in night-migratory songbirds [27,29,30]. These results exclude previous suggestions that a radical pair involving a counter-radical with very few or no hyperfine interactions could be responsible for magnetoreception in night-migratory songbirds [25–27,30,70]. In short, broadband RF fields are much more likely to affect a biological radical pair reaction than any particular single-frequency field and the upper and lower cut-off frequencies can provide hints as to the nature of the radicals involved.

Quantum chemical calculations based on the FAD–Trp radical pair and its hyperfine interactions in *Er*Cry4a predict the frequency range of broadband RF fields which should disrupt the avian magnetic compass [29,30,70]. According to these predictions, an FAD-containing radical pair should be affected by broadband RF fields up to 100–120 MHz [29,70]. Leberecht et al. [30] showed that a 75–85 MHz RF field disrupts the avian magnetic compass, which demonstrated the involvement of at least one radical with hyperfine interactions similar in strength to those in FAD [70]. These spin dynamics calculations also predicted that, in the frequency range 120–220 MHz, there would be a negligibly small probability of disrupting the avian compass, while at any frequency beyond 220 MHz, there should be no possibility of disruption [29,70]. Magnetic compass orientation experiments performed in the presence of RF noise in the ranges 140–150 MHz and 235–245 MHz showed that neither of these broadband fields affected the avian magnetic compass in behavioural tests of night-migratory songbirds [29].

As mentioned above, different radicals have different hyperfine interactions and the maximum radiofrequencies expected to affect different radical pairs can be calculated [29,70]. For TrpH^•+^ and FAD^•−^ radicals in *Er*Cry4a, these maximum frequencies are 106 MHz and ∼116 MHz, respectively. For a radical pair, the upper cut-off frequency is mostly determined by the larger of the two maxima, in this case ∼116 MHz (for FAD^•−^in *Er*Cry4a [29]). Broadband RF fields beyond the predicted ∼116 MHz cut-off were estimated to have a low total probability (∼1%) of affecting the radical pair reaction. Such small effects are highly unlikely to be detectable in behavioural experiments [29]. Furthermore, the likelihood of a broadband RF field having a disruptive effect on the avian magnetic compass decreases as the frequency of the broadband field approaches the cut-off from below. Potential consequences for the conservation of migratory songbirds make it relevant to test whether RF fields near the predicted cut-off frequency can disrupt their magnetic compass orientation. The aim of the present paper therefore is to test the magnetic compass orientation behaviour of a night-migratory bird under a broadband RF field approximately centred around the predicted upper cut-off frequency for a FAD–Trp radical in *Er*Cry4a (i.e., 110–120 MHz).

## 2 Material and Methods

### 2.1 Test animals

The procedures for the behavioural tests were identical to those of our two recent studies testing broadband RF fields [29,30]. Full details can be found in those papers and their Supplementary Information Appendices. Here we will provide a short summary. The major difference to Leberecht et al. [29,30] was the chosen RF frequency-band (110–120 MHz). We tested Eurasian blackcaps (*Sylvia atricapilla*) in the spring migratory seasons of 2022 and 2023. The birds had been wild-caught during autumn migration in the immediate vicinity of the University of Oldenburg. When possible, the birds were kept in pairs in on-site cages in a windowless room under a light regime imitating the local photoperiod, with *ad libitum* access to food and water. The onset of their spontaneous, nocturnal migratory restlessness activity was surveyed by infrared-based activity loggers installed in each cage.

### 2.2 Generation and measurement of static magnetic field stimuli

Behavioural experiments were performed in two static magnetic field conditions, the normal magnetic field (NMF) and a changed magnetic field (CMF). In the NMF condition the local geomagnetic field of Oldenburg was present. In the CMF condition, the horizontal component (declination) of the magnetic field was rotated 120° counter-clockwise while keeping the local magnetic field strength and inclination angle unchanged [4,5,26–30]. The behavioural experiments were conducted on a wooden table in the centre of the coil system where the homogeneity of the applied static magnetic field was at least 99%. Each of the three sets of four coils was powered by a separate constant-current power supply (BOP 50–4 M, Kepco Inc., Flushing, NY, USA). In each chamber, the local and the 120°-counter-clockwise rotated magnetic fields were recorded daily at alternating, opposite corners and in the centre of the experimental table using a flux-gate magnetometer (FVM-400, Meda Inc., Dulles, VA, USA).

In both the NMF and CMF conditions, the same amount of current ran through a double-wrapped, three-axis Merritt four-coil system (ca. 2 m × 2 m × 2 m) in each of the three chambers [4,5,26–30]. The NMF condition was generated by running currents anti-parallel through the two double-wrapped sets of windings. Because the magnetic fields induced by currents running in opposite directions through the two identical sets of windings in each coil exactly cancel one another, the birds perceived only the local geomagnetic field. In the CMF condition, the currents ran in parallel through all the windings of the double-wrapped coils. The average magnetic measurements of the NMF over all test days and chambers were: inclination 67.51° ± 0.2° (mean ± standard deviation (s.d.) for circular data) at an average intensity 48,843 ± 259 nT. In the CMF condition, the average inclination was 67.63° ± 0.26°, with an average horizontal direction (declination) of –120° ± 0.89° and an average intensity of 48,845 ± 300 nT.

### 2.3 Generation and measurement of time-dependent electromagnetic fields

The behavioural experiments took place in a specially constructed, non-magnetic laboratory building described in earlier studies [28–30, full description in 26]. Within the laboratory, three aluminium-shielded chambers acting as Faraday cages allowed static magnetic fields to pass through, while attenuating time-dependent electromagnetic fields, ranging from 10 kHz to 10 GHz, by a factor of at least 10^5^. The electrical equipment used to generate the RF fields was grounded through an 8 m-deep earthing rod, while the individual chambers were grounded using single electrode loops in the laboratory base [26].

The RF fields were produced with one signal generator per experimental chamber (Rhode and Schwarz, SMBV100A (9 kHz–6 GHz) and SMBV100B (8 kHz–3 GHz), Germany), set to generate broadband noise in the spectral range 110–120 MHz (centre frequency: 115 ± 5.75 MHz, with a stimulus plateau width of 10 MHz (110–120 MHz)). The experimental groups of birds that experienced this RF-stimulation are denoted with a “-RF” suffix. In the RF-exposure condition, the output RMS voltage at the signal generator was set to 37 mV, while in the control condition, the RMS voltage was set to 15 nV, so that in both conditions, the signal generators were actively producing a signal. Unless noted otherwise, coaxial cables (Schwarzbeck Mess-Elektronik, AK 9515 E, 50 Ohm, N-Connector, Germany) were used.

The signals were fed into broadband power amplifiers (AR Deutschland GmbH, 50U1000 (10 kHz–1000 MHz), Germany), amplified to 53% of the maximum gain (ca. 45 dB) and guided into the experimental chambers via a wall panel. Inside the chambers, the signal was passed to a custom-built bandpass filter box (attenuation up to 71.88 MHz; 3 dB attenuation at 71.88 MHz), followed by an 8-Way splitter (Werlatone, Model D5829-10, 20–500 MHz, 400 W, N-Connectors, Patterson, NY, USA). Both were located under the experimental table within the Merritt four-coil system. Each of the eight outputs from the splitter was fed into a coaxial cable (RG58C/U MIL-C-17, BNC connector), which was wrapped in a single turn around a custom-built, circular PVC antenna frame (diameter: 35.7 cm, height: 9 cm, circumference: 112.2 cm). The shield of the coaxial cable was removed along the circumference of the single turn coil, and the inner conductor was connected at one end to the shield at the other end to close the loop. This single-loop magnetic coil acted as an emittance antenna, applying the generated broadband noise to the Emlen funnels placed inside the antennas. The generators were in ‘RF OFF’-mode and only switched to ‘RF ON’-mode for the experiments or during measurements.

Every day, after the daily measurement of the static magnetic field, the magnetic components of the RF fields were measured in each chamber at opposite corners or edges of the tables. As recommended by Hiscock et al. [70] and carried out in Leberecht et al. [29,30], the fields were measured with a calibrated active loop antenna (Schwarzbeck Mess-Elektronik, HFS 1546, 150 kHz–400 MHz, Germany), placed 1.5 cm above the centre of the emission antennas. The measurement antenna was connected through the wall panel to a signal analyser (Rhode and Schwarz, FSV3004 Signal and Spectrum Analyser (10 Hz–4 GHz), Germany). The electric components of the fields were measured similarly with a calibrated active bi-conical antenna (Schwarzbeck Mess-Elektronik, EFS 9218 (9 kHz–300 MHz), Germany). The RF fields were recorded for 1 min daily with the resolution bandwidth of the analyser set to 10 kHz. Before and after a migratory season, a one-hour measurement took place to record the RF fields applied during the experimental phase (magnetic component: figure 1; electric component: see SI-figure 1). For comparability with earlier studies [26–30], we recorded two different spectral traces in each measurement: ‘maxhold’ (with the “Maxpeak” detector; for the detection of absolute maxima at a given frequency) and average spectral amplitudes (with the “RMS” detector; measuring the root-mean-square intensity over time at a given frequency). The spectral intensity of the applied broadband noise in the 110-120 MHz frequency range (intensity: 2022: *B*_rms_ = 2.1 nT, 2023: *B*_rms_ = 1.8 nT; density: 2022: *b*^max^ = 2.16 pT/√Hz, 2023: *b*^max^ = 2.24 pT/√Hz) integrated according to the equations in Kobylkov et al. [28] was comparable to previous studies (see SI-table 1). All measurements fall between the respective values of the broadband RF fields causing disorientation in European robins [27] and Eurasian blackcaps [30]. We actually tried to apply stronger RF intensity in the 110-120 MHz range, but at higher amplification, unwanted side bands started to appear, so the used intensities were the strongest possible “clean” 110-120 MHz stimulus with our equipment for the given frequency range.

**Figure 1:**
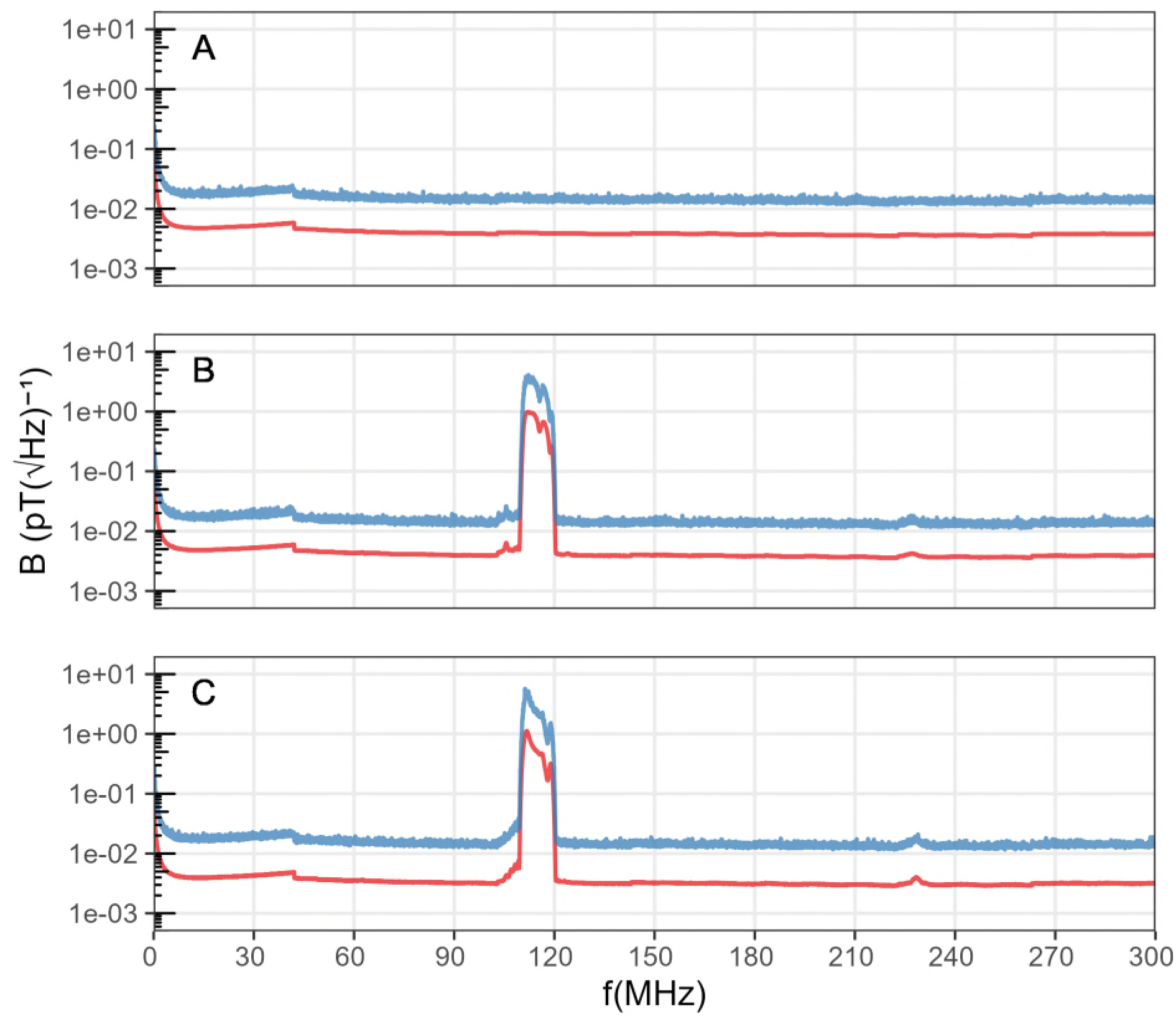
Measurements of the magnetic components of the control condition. (A), the 110–120 MHz broadband RF field conditions in 2022 (B), and 2023 (C) across the range from 150 kHz to 300 MHz. Spectral traces: ‘average’ (lower red line); ‘maxhold’ (upper blue line).

### 2.4 Acquisition and analysis of behavioural data

As soon as Eurasian blackcaps had been pre-screened by testing each individual’s migratory motivation and ability to orient using their magnetic compass in the NMF and CMF conditions without RF-noise, they were re-tested in the control and RF-exposure conditions (2022: controls: 23 March to 4 May; RF condition: 21 April to 3 May; 2023: controls: 31 March to 11 May; RF condition: 8 April to 10 May). The pre-screening results are not shown in figure 2, as these data were exclusively used for selection of motivated birds and therefore excluded from further use. The behavioural orientation data in the NMF and CMF conditions with and without RF-noise are reported in figure 2. On every test day, the birds were caught from their housing cages one hour before the end of civil twilight (approximately 30 min before sunset) to allow them to experience the sunset and potentially to calibrate their magnetic compass [77,but see 78]. The birds were placed in transport boxes with a net-covered opening facing the sunset in the nearby botanical garden and sheltered from rain and wind. During the transfer to the non-magnetic experimental laboratory, the transport boxes were covered with a black cloth to exclude any disturbances from streetlights. As in previous studies [4,5,26–30], the behavioural tests were conducted under dim light (mean ± s.d. = 2.42 ± 0.05 mW m^-2^) produced by incandescent light bulbs.

**Figure 2:**
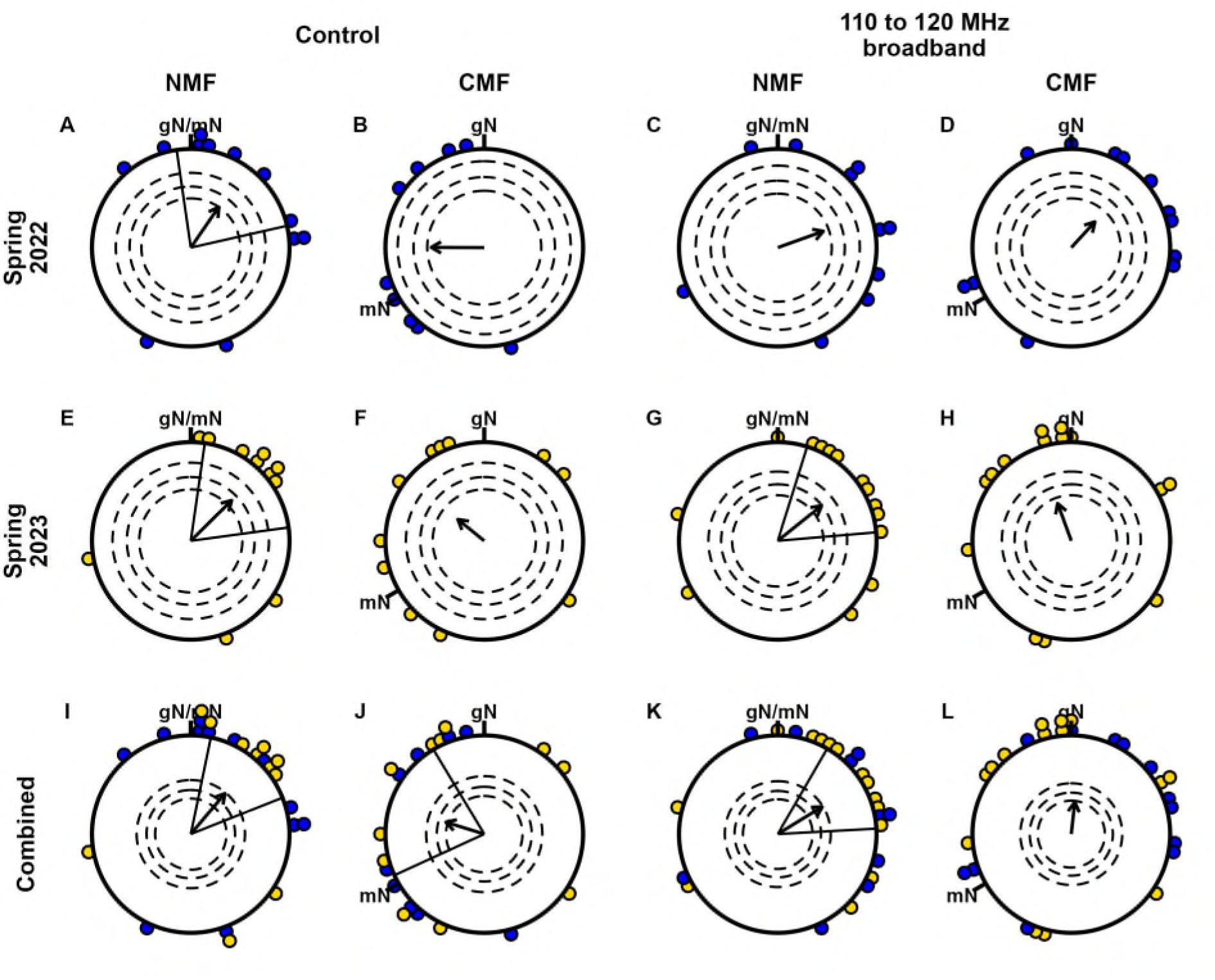
Magnetic compass orientation of Eurasian blackcaps. Rows: The magnetic compass orientation in the spring migratory seasons of 2022, 2023, and their combination. Columns: Magnetic compass orientation under different magnetic field conditions in the absence (Control) and presence of 110–120 MHz RF fields: NMF — normal magnetic field in Oldenburg; CMF — 120°-CCW-rotated magnetic field (only the magnetic declination changed). Each coloured dot represents the mean direction of one individual bird rounded to the nearest 5°. The arrows display the group mean orientation and the arrow length is calculated by adding up unit vectors for each of the individual birds’ mean direction and dividing by the number of tested birds. The dashed circles indicate the radii of the group mean vectors needed for directional significance according to the Rayleigh test (inner: p < 0.05, middle: p < 0.01, outer: p < 0.001). Radial lines flanking the group mean vector mark the 95% confidence interval for the group mean direction. gN: geographical North; mN: magnetic North. Blue dots: birds from the 2022 cohort; yellow dots: birds from the 2023 cohort. Although the same individuals were tested in all 4 conditions, the sample size is slightly different between the conditions, since in some cases, a particularbird did not provide data in all tested conditions (see Section 2.4 ‘Acquisition and analysis of behavioural data’).

The behavioural experiments were performed using modified Emlen funnels made of white PVC (35 cm diameter, 15 cm high, walls 45° inclined; [79,80]). During the migratory season, night-migratory songbirds exhibit “migratory restlessness”, expressed as jumping around and wing-whirring in a cage. Birds displaying migratory restlessness at night tend to jump in the direction they want to migrate. The sloping sides of the funnels cause them to slide back down leaving marks on the scratch-sensitive thermal paper with which the funnels are lined [80], allowing the birds’ intended migratory directions to be recorded. The edges of the scratch papers were joined with adhesive tape and the resulting overlaps were aligned with one of the cardinal directions, chosen at random for each experiment and experimental chamber.

In the experimental chambers, each bird was placed in a designated Emlen funnel lined with scratch-sensitive thermal paper. After one hour, the birds were returned to their transport boxes. Every test day consisted of two to three rounds of tests in the same experimental condition with only the position of the funnel on the experimental table changed between rounds for each bird. After the last experimental round, the birds were returned to their housing cages. After every experimental round, the scratch papers were collected, the funnels cleaned and, if applicable, prepared for another round. The behavioural tests were not blinded: the researcher who put the birds into the funnels knew the conditions under which they would be tested. However, the scratch papers were analysed independently and in a blind manner by two researchers, unaware of the static magnetic field and RF field conditions and the geographical alignment of the scratch papers. Papers that had fewer than 30 scratches (143 of 988 papers, 14.5%) were classified as inactive and excluded from further analysis. For each of the remaining papers, the mean orientation of the scratches was determined to the nearest 10°. If the orientation directions estimated independently by the two researchers agreed, i.e. were within 30° of each other, the mean of the two values was taken as the bird’s orientation (767 of 988 papers, 77.6%). Otherwise, the paper was reassessed by a third evaluator and if the third direction also differed by more than 30° from either of the first two evaluations, the paper was deemed to be random (78 of 988 papers, 7.9%).

The individual orientation results for every bird in each condition were added up as unit vectors and divided by the number of test results to calculate each of the individual bird mean vectors, following Batschelet [81]. The direction of the individual mean resultant vector represents the mean direction of a bird in a given condition. The length of the individual mean vector represents how consistently a bird oriented in the same direction in a given condition (the individual *r*-value, or so-called directedness). The vector calculations were performed with a custom-written R-script using circular statistics [82–84]. The data used for the final analysis are accessible in SI-table 2. In line with previous studies [5,28–30], in figure 2 we included the means of all individuals with at least 3 directed tests in the relevant condition and an individual mean vector *r* ≥ 0.2. We also required that any given individual bird contributed a directional value to at least two of the four test conditions. For every experimental condition, the group mean orientation and directedness were then calculated by adding up unit vectors in the mean orientation direction of each bird and dividing by the number of birds tested. The group mean orientation was tested against the null hypothesis of a uniform circular distribution (Rayleigh test) and a 95% confidence interval around the mean direction was calculated. For the comparison of the directions between the test conditions, the Mardia-Watson-Wheeler test was used [81] and the group rotation of the orientation between significant NMF and CMF conditions was tested with the V-Test [81, as recommended by Landler et al. 85].

To investigate the potential disorienting effect of our experimental broadband 110–120 MHz RF-field, we used the Levene Test [86, applied for orientation data in 29,87] to test the equality of variance between conditions, a bootstrapping approach [23,29,30,88,89] to test whether the directions in the CMF-RF condition were significantly more spread out (did they show a significantly lower *r*-value?) than the CMF control condition [23,29,30,88,89], and a randomization approach to compare the group directedness with previous studies testing broadband RF fields in spring [28–30]. The bootstrapping procedure and randomization approach are described in detail in the supplementary information (SI-C Supplementary analysis).

## 3 Results

Similar to free-flying conspecifics [90], the group of birds tested in the normal magnetic field of Oldenburg (NMF) displayed a significant, seasonally appropriate orientation towards NE in both years (Rayleigh Test: *N* = 23, mean ± s.d. = 40° ± 55°, 95% CI = 12° – 68°, *r* = 0.54, *Z* = 6.65, *p* = 0.0009; figure 2*I*). When the magnetic field was turned 120° counter-clockwise (CCW; CMF condition), the birds adjusted their group heading accordingly with a 113° CCW turn, orientating significantly towards geographic W and magnetic NE (*N* = 20, mean ± s.d. = 287° ± 62°, 95% CI = 245° – 329°, *r* = 0.41, *Z* = 3.3, *p* = 0.0348; figure 2*J*). The directions in the NMF and CMF control conditions were significantly different from each other (Mardia-Watson-Wheeler (MWW): *W_(degrees of freedom = 2)_* = 10.75, *p* = 0.0046). The compensation for the 120° CCW turn of the magnetic field was also statistically significant (*V* = 0.4, *µ* = 2.55, *p* = 0.005, V-test of CMF mean orientation data against the group mean data of the NMF condition (adjusted for the 120°-CCW horizontal magnetic field rotation)).

### 3.1 Effects of a 110–120 MHz broadband RF fields

In the NMF condition with 110–120 MHz broadband RF fields present (NMF-RF), the birds’ significant orientation towards the seasonally appropriate NE direction (*N* = 24, mean ± s.d. = 58° ± 56°, 95% CI = 30° – 87°, *r* = 0.53, *Z* = 6.72, *p* = 0.0008; figure 2*K*) did not differ from the control condition (MWW: *W_(2)_* = 4.25, *p* = 0.1192). With the magnetic field turned (CMF-RF), the birds displayed a non-significant group orientation towards geographic North (*N* = 26, mean ± s.d. = 6° ± 66°, *r* = 0.33, *Z* = 2.78, *p* = 0.0606; figure 2*L*). The mean orientation of the birds’ is rotated 52° CCW compared to the NMF condition, but this difference in orientation was not statistically significant (*V* = 0.12, *µ* = 0.89, *p* = 0.1878) and the CMF-RF data were also not statistically different from either the NMF-RF, or the control CMF condition (SI-C table 4).

We did not detect any significant differences in the variance of the individual directedness *r*-values between the test conditions (Levene Test – *F _d.f. (condition, residuals)_*: *F_(3,89)_* = 1.215, *p* = 0.309). According to the results of the bootstrap analysis (see SI-C Supplementary analysis), the bootstrapped *r*-value and angular 95% confidence intervals (CI) of the CMF and CMF-RF condition overlapped (SI-C table 7). Furthermore, the observed group directedness lay within the *r*-value CI of the respective other condition (*r*_CMF-RF_ = 0.327, *r*-CI_CMF-RF_ = 0.105 – 0.627; *r*_CMF_ = 0.406, *r*-CI_CMF_ = 0.213 – 0.63). Many of the bootstrap iterations of the CMF conditions were at least as directed as the CMF-RF condition and vice versa (CMF: 83.6%; CMF-RF: 38.7%). However, very few iterations also fell in the angular CI of the respective other condition (CMF: 3.5%; CMF-RF: 1.7%). Thus, the CMF conditions with and without broadband RF fields can be considered as significantly different in their orientation, although similar in their directedness (SI-C table 4, 7 and SI-C figure 3).

To compare the orientation of our birds in the CMF-RF condition under broadband 110–120 MHz RF fields with previous spring migratory seasons under other broadband RF fields [28–30] we employed a randomization approach (see SI-C Supplementary analysis). The directedness of the present CMF-RF data lay in-between random and oriented samples of previous studies with its 95% CIs for *r*_group_ overlapping with both random and oriented previous samples. Thus, the orientation of the birds tested in the CMF under broadband 110–120 MHz RF fields is not significantly different from either the previously reported oriented or disoriented groups (figure 3 and SI-C table 8).

**Figure 3:**
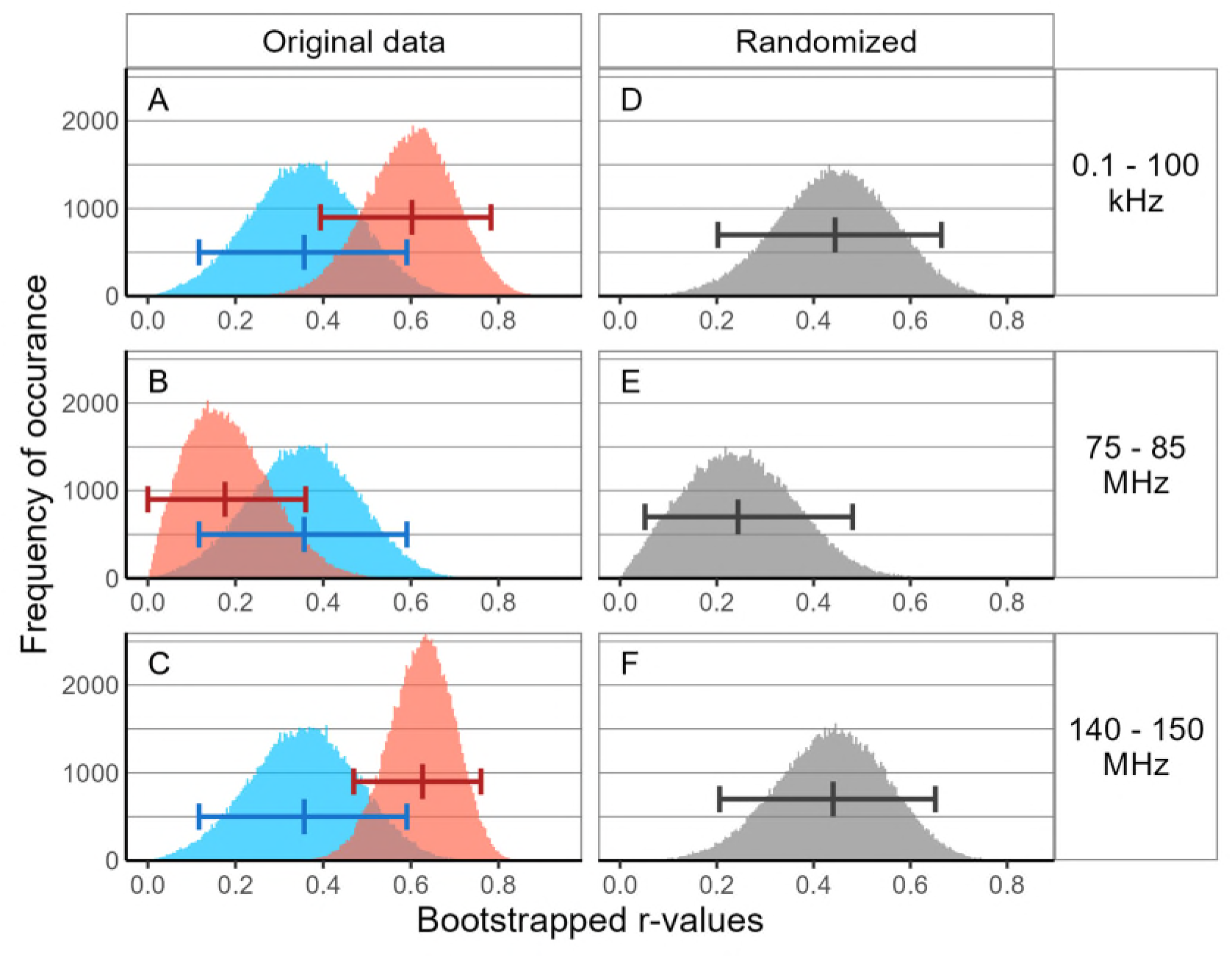
A-C: Histograms of the bootstrapped r-values of the original data in the CMF condition in the presence of broadband RF fields (CMF-RF), in the present study from 110–120 MHz (blue) beside the CMF-RF condition of previous studies (red). D-F: Histograms of the r-values from the randomized sampling from the combination of the CMF-RF data of the present study and the respective previous studies. (A, D) Kobylkov et al. 2019 [28] with non-disruptive broadband fields from 0.1–100 kHz, (B, E) Leberecht et al. 2022 [30] with disruptive broadband fields from 75–85 MHz, and (C, F) Leberecht et al. 2023 [29] with non-disruptive broadband fields from 140–150 MHz. The CMF-RF data from the present study under 110-120 MHz RF fields (blue histograms) were combined across years and 26 data points (the sample size of the present CMF-RF condition) were randomly sampled with replacement. The process was repeated 100,000 times and for each r-value was computed. The whiskers indicate the median and 95% confidence intervals of the r-value. Number of bins = 200.

### 3.2 Differences between experimental seasons

In both years (2022 and 2023), the NMF control conditions were significantly oriented in the seasonally appropriate migratory direction of NE (figure 2*A* and *E*; SI-C table 3). In the CMF condition, the groups of birds in both years tended non-significantly towards W and NW, respectively (roughly magnetic NE; figure 2*B* and *F*; SI-C table 3). Neither the control NMF, nor CMF were statistically different between the years (NMF: *W_(2)_* = 0.78, *p* = 0.677; CMF: *W_(2)_* = 2.85, *p* = 0.2408).

In the presence of 110–120 MHZ broadband RF fields of intensity *B*_rms_ = 2.1 nT, the group of birds of the 2022 cohort displayed non-significant tendencies towards NE, both in the NMF-RF and CMF-RF condition (figure 2*C* and *D*; see SI-C table 3). The lack of significant group orientations and lack of a significant turn between NMF-RF and CMF-RF could indicate a disorienting effect of the 110–120 MHz broadband RF fields (SI-C table 3–5).

In the NMF-RF condition of 2023, with RF fields of intensity *B*_rms_ = 1.8 nT, the birds oriented significantly in the seasonally appropriate NE migratory direction (*N* = 14, mean ± s.d. = 51° ± 53°, 95% CI = 17° – 85°, *r* = 0.57, *Z* = 4.52, *p* = 0.0085; figure 2*G*), and tended non-significantly towards geographic NNW (magnetic East) in the CMF-RF condition (*N* = 14, mean ± s.d. = 340° ± 63°, *r* = 0.4, *Z* = 2.29, *p* = 0.1001; figure 2*H*). While the NMF-RF and CMF-RF conditions in 2023 were statistically different from each other, neither differed from its respective control and the turn of 71° between them was not statistically significant (SI-C table 3–5).

As the group mean orientations in the NMF-RF condition of both years were directed towards NE, regardless of significance, there was no statistical difference between them (*W_(2)_* = 0.58, *p* = 0.7487). The same was found for the comparison of the CMF-RF condition between the years (*W_(2)_* = 1.48, *p* = 0.4769). In general, the variances of individual *r*-values between the test conditions did not differ significantly across the years (Levene Test: *F_(7,85)_* = 1.01, *p* = 0.4305), and we also did not find a statistical difference in the variance of the birds’ orientation angles (SI-C table 6). Therefore, neither year, nor test condition had a statistically significant effect on the spread of the birds’ orientations or the individual’s consistency in orientation. A more detailed comparison of the experimental seasons can be found in SI-C Supplementary Analysis.

### 3.3 Result summary

Without broadband RF fields present, we observed the expected magnetic compass orientation behaviour. We found potential RF effects in the CMF-RF condition, but not in the NMF-RF condition. Differences in the condition-specific direction of groups across the years only occurred in the presence of the RF field. It could not be determined whether this effect across the years is related to the presence of the RF field or its intensity, as there was no difference in the variance from the condition-specific group direction or directedness. From our bootstrap analysis of the *r*-values, we can conclude that the birds as a group oriented equally well in the condition-specific group direction, whether RF fields were present or not. Our randomization analysis suggests that our present RF data fall between the observation of oriented and disoriented birds from previous studies. We discuss the interpretation of these results in the context of the presence of the RF fields and their intensity in the following.

## 4 Discussion

Previous studies provided a clear interpretation of the effect of broadband RF fields on the avian magnetic compass orientation behaviour during two experimental migratory seasons [26–30]. Our experiments under broadband RF fields from 110–120 MHz did not yield a similarly clear outcome. The lack of an appropriately rotated orientation in the turned magnetic field (CMF) in the presence of our broadband RF field could be interpreted as evidence for a disruptive effect, but such a conclusion should be treated with caution. Under broadband 110–120 MHz RF fields, the birds oriented correctly in the appropriate migratory direction in the NMF-RF condition, while in the CMF-RF condition, neither clear orientation, nor clear disorientation was observed.

### 4.1 Potential disorienting effects of a 110–120 MHz broadband RF field

Our experimental broadband 110–120 MHz RF field included the predicted cut-off frequency, ∼116 MHz, for the avian magnetic compass sensor, where the probability to disturb the avian magnetic compass was predicted to be smaller than at lower frequencies [29,70]. In contrast to the previous binary predictions of the RF field effects on the orientation of birds (oriented: [29,70]; disoriented: [30,70]), there is no clear oriented-disoriented-prediction for the RF region around the predicted cut-off frequency.

As our tests under RF fields and controls were always interleaved (sometimes on the same day), an effect of the progression of the migratory season is unlikely, as it would also have affected the controls, and no such effect was observed in the control. Given that the group orientations under control conditions in both present and past studies reflect the expected rotation of the orientation to a turned magnetic field under control conditions throughout the migratory seasons [1,4,5,26,27,29,30], we find progression-of-season-effects extremely unlikely.

Curiously, our CMF-RF behavioural data seem not to be as randomly distributed as previous studies that demonstrated disruptive effects of broadband RF fields [25–27,30]. According to our bootstrap and randomization analyses (figure 3 and see SI-C table 8), it seems that the orientation in our CMF-RF condition falls in-between the orientation observed in previous studies investigating RF field ranges with no disruptive effect [28,29] and the lack of orientation observed under disruptive broadband RF fields [30].

One reason for this ambiguous result could be that the RF fields from 110–120 MHz at the intensities tested were not fully disruptive. Our RF field intensities and noise densities fell in-between the values of two prior studies showing disorienting effects on night-migratory songbirds (see SI-table 1; [27,30]), hence our RF fields should have been strong enough to evoke similar effects. The aforementioned gradually decreasing probability for an RF effect close to the predicted cut-off [29,70] could therefore be reflected in the unclear behavioural response we observed. Thus, it is in fact not so surprising, that the orientation performance of our birds tested under 110–120 MHz fields seems to fall in a “grey zone” inconsistent with a binary “oriented–disoriented” interpretation.

Although purely speculative, the ambiguity arising from the behavioural data (figure 2) and the inconclusive categorisation (figure 3) of the behaviour under 110–120 MHz fields into oriented or disoriented could be an indication that the birds’ magnetic compass was partly but not fully disrupted. This could be because FAD^•−^, which has a predicted upper cut-off frequency of ∼116 MHz [29,70], might have been affected, whereas TrpH^•+^ was most likely not affected, since its predicted cut-off (106 MHz) lies below the broadband RF range we tested here. The likelihood of a disruptive effect below ∼116 MHz within our 110–120 MHz broadband RF field is predicted to be low [29,70], but could still occasionally have an effect on the birds orientation. We currently have no explanation for why the suspected partly disruptive effect seemed to occur more strongly in the CMF-RF condition than in the NMF-RF condition.

We did ask ourselves whether any possible effect could come from the direction of the RF magnetic field with respect to the static field. The prediction from radical pair theory ([20], specifically Section 3.1 in its Supporting Information) is that there would only be a directional dependence if (a) one radical had no hyperfine interactions and (b) the dipolar interaction was effectively zero. Neither condition is at all likely (for reference see [9,91]). Furthermore, the experimental setup and thus the orientation of the RF fields were identical to both of our previous studies around 80 and 145 MHz [29,30] and the RF-field orientation was identical in the NMF and CMF conditions. We therefore find it extremely unlikely that any RF-orientation effect could have played a role in these experiments. Finally, we would like to mention that it has been suggested that the disorienting effects of RF noise could imply the existence of a specialised magnetite-based detector whose function is to “switch off” the magnetic compass during terrestrial and/or solar storms when magnetic cues might be less reliable [23,92,93]. Leaving aside the absence of both supporting evidence and a plausible sensing mechanism for such an exquisitely sensitive receptor, it would be a rather remarkable coincidence if such a sensor would have a cut-off frequency very close to that predicted by the magnetic properties of radical pairs in cryptochrome.

### 4.2 Annual effects

We do not think that the minor difference in RF field intensity between the years is a main driver behind the ambiguous results in the CMF condition. For single frequency RF fields, the existence of a RF intensity threshold for the disruption of the avian magnetic compass has been reported [20,22]. However, broadband RF fields are known to require much lower intensities to cause disorientation in the birds [25–27,30] and no specific intensity threshold is currently documented. As stated in the previous section (and the Methods section), the RF field strengths used in the present study fell well within the range of previous studies showing disorienting effects of broadband RF fields (see SI-table 1). Furthermore, the group directedness fell between previous studies with disorienting/not disorienting broadband RF fields (see SI-figure 4 and SI-C table 8). While we do not doubt the existence of a RF intensity threshold, we deem it unlikely that our RF field intensities, by chance, were set to intensities right around such a threshold.

### 4.3 Future prospects

Establishing an intensity threshold for disruptive effects of broadband RF fields by means of behavioural experiments would require an exhaustive brute-force approach [29,30], as behavioural experiments are season-dependent and prone to noisy data. Thus, it is not feasible to use behavioural experiments to resolve whether the ambiguous effects of our 110–120 MHz RF field arise from an intensity threshold or because the broadband RF field spanned the predicted cut-off frequency at ∼116 MHz or from a combination of both of these potential reasons [29,70].

Leberecht et al. [30] noted in their discussion section that behavioural experiments may not possess the necessary resolution to exactly confirm the upper cut-off frequency for the radical pair underlying avian magnetoreception. Our results may reflect this suboptimal resolution of behavioural experiments. We agree with Leberecht et al. [30] that further exhaustive behavioural tests are most likely not going to give a clearer picture of the maximum frequency at which RF fields can disrupt the magnetic compass sense. Instead, an effort should be made to establish a reliable and reproducible electro-physiological readout from the avian retina. In principle, recordings in different magnetic fields should provide different signals. Indeed, electroretinographic responses to changes in the magnetic field have been reported [94]. However, these differences were minute and have not been broadly reproduced yet. Once consistent, independently reproducible neuronal responses to different magnetic fields are established, precisely controlled RF fields can be introduced. The stable neuronal response from the retina to a magnetic field should change, once broadband RF fields start to disrupt the radical-pair mechanism. With such an electrophysiological approach, the RF spectrum just above and below the cut-off frequency of ∼116 MHz and potential RF intensity thresholds can be probed. To avoid artefacts, it is crucial that electrophysiological recordings of magnetically neural signals from the avian retina are performed in a meticulously controlled non-magnetic environment and that artefacts that can occur due to interference between the changes of the magnetic field and the necessary electronic devices are carefully considered.

## Ethics

All experimental procedures were conducted in accordance with national and local guidelines for the use of animals in research, approved by the Animal Care and Use Committees of the Niedersächsisches Landesamt für Verbraucherschutz und Lebensmittelsicherheit (LAVES, Oldenburg, Germany, 33.19-42502-04-17/2724).

## Data, Materials, and Software Availability

Code is available on request. All study data are included in the article and/or SI Appendix.

## Acknowledgements

The authors gratefully acknowledge the University of Oldenburg’s workshops for expert technical assistance and its animal keeping facility and veterinarians for taking care of our birds.

## Author contributions

HM and PJH designed the research; BL, BS, LS, and LV performed the experiments; BS, LB, JO and FI evaluated the blinded data; BL performed the statistical analysis; JW provided intellectual input to the statistical analysis and interpretation; HM provided reagents and facilities. The first draft was written by BL and further improved by BS, JW, HM and PJH. All authors read and commented on the manuscript.

