## Supplementary Information for "Magnetic compass orientation behaviour of Eurasian blackcaps at the predicted 110–120 MHz upper cut-off frequency for radiofrequency-field effects"

ORCID numbers:

Bo Leberecht: 0000-0001-9672-1504

Baladev Satish: 0009-0009-0086-6094

Leonard Schwigon: 0009-0004-1453-2638

Lalitha Venkatraman: 0000-0003-4995-5652

Joe Wynn: 0000-0002-5552-6435

P. J. Hore: 0000-0002-8863-570X

Henrik Mouritsen: 0000-0001-7082-4839

### A Supplementary figures

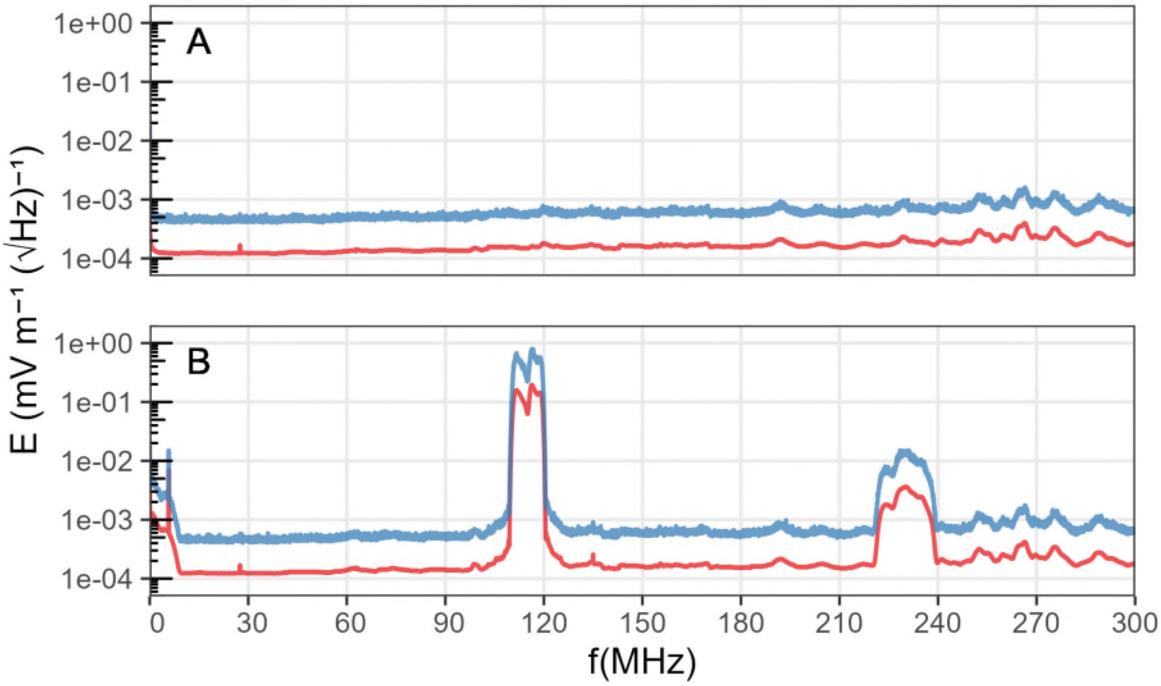


Figure 1: *Measurements of the electric components of the control condition (A) and the 110–120 MHz broadband RF field conditions (B) across the range from 150 kHz to 300 MHz. Spectral traces: ‘average’ (lower red line); ‘maxhold’ (upper blue line). Note that the otherwise clean electrical spectrum of (B) shows some weak harmonics around 230 MHz..*

#### 43 B Supplementary tables

*SI-table 1: Comparative table of different noise density and intensity measures from Kobylkov et al. (2019) and Leberecht et al. (2022, 2023) and estimates from Engels et al. (2014), with the results of the present study appended. Species: ER: European robin, BC: Eurasian blackcap. For formulas for* $\bar{b}^{max}$, $\bar{b}$, $B^{max}_{rms}$, $B_{rms}$ *see Eqs (2.1) and (2.3) in Kobylkov et al. (2019). Orientation: behavioural experiments resulted in oriented (+), disoriented (−) or ambiguously oriented (?) birds.*

| Frequency band (kHz) | Species | $\bar{b}^{max}$ (pT/√Hz) | $\bar{b}$ (pT/√Hz) | $B^{max}_{rms}$ (nT) | $B_{rms}$ (nT) | Orientation | Source |
|---|---|---|---|---|---|---|---|
| 10 − 5000 | ER | 0.07 | ~0.02 | 1.18 | ~0.37 | + | Engels et al. 2014 (Fig.4f, blue trace) |
| 10 − 5000 | ER | 5.58 | ~1.8 | 23.2 | ~7.35 | − | Engels et al. 2014 (Fig.4f, red trace) |
| 20 − 450 | ER | 30.1 | ~9.5 | 23.1 | ~7.29 | − | Engels et al. 2014 (Fig.4f, green trace) |
| 600 − 3000 | ER | 1.35 | ~0.43 | 2.3 | ~0.73 | − | Engels et al. 2014 (Fig.4f, black trace) |
| 0.1 − 100 | BC | 23.3 | 6.6 | 7.98 | 2.26 | + | Kobylkov et al. 2019 (Fig.1a, RF−on) |
| 0.1 − 100 | BC | 0.92 | 0.32 | 1.29 | 0.51 | + | Kobylkov et al. 2019 (Fig.1c, RF−off) |
| 75,000 − 85,000 | BC | 0.015 | 0.004 | 0.046 | 0.013 | + | Leberecht et al. 2022 (Fig.2A, RF−off) |
| 75,000 – 85,000 | BC | 2.525 | 0.701 | 8.681 | 2.404 | − | Leberecht et al. 2022 (Fig.2B, RF−on) |
| 235,000 – 245,000 | BC | 0.014 | 0.004 | 0.044 | 0.012 | + | Leberecht et al. 2023 (Fig.S5A, RF−off) |
| 235,000 – 245,000 | BC | 3.601 | 0.961 | 12.359 | 3.286 | + | Leberecht et al. 2023 (Fig.S5B, RF−on) |
| 140,000 – 150,000 | BC | 0.012 | 0.004 | 0.037 | 0.012 | + | Leberecht et al. 2023 (Fig.S5C, RF−off) |
| 140,000 – 150,000 | BC | 2.813 | 0.948 | 9.463 | 3.181 | + | Leberecht et al. 2023 (Fig.S5D, RF−on) |
| 110,000 – 120,000 | BC | 0.015 | 0.004 | 0.046 | 0.012 | + | Present Study (Fig.1A, RF−off) |
| 110,000 – 120,000 | BC | 2.161 | 0.603 | 7.568 | 2.108 | (−/?) | Present Study (Fig.1B, RF−on, 2022) |
| 110,000 – 120,000 | BC | 2.236 | 0.501 | 8.056 | 1.808 | (+/?) | Present Study (Fig.1C, RF−on, 2023) |

*SI-table 2.Experimental result summary for the individual birds in the respective test conditions (NMF: normal magnetic field in Oldenburg; CMF: changed magnetic field, turned by 120° counter-clockwise; NMF-RF: NMF with 110–120 MHz broadband RF fields present; CMF-RF: CMF with 110–120 MHz broadband RF fields present). For each individual (Ring) the mean orientation (Angle), Rayleigh value (r-value) for this orientation, the portions of oriented, random and inactive trials for the overall number of trials (N) are listed. Result with insufficient valid trials or undirected orientation (valid < 3; r < 0.2) are marked in red and were not used for the final orientation statistics and diagrams.*

| | | NMF | | | | | | CMF | | | | | | NMF-RF | | | | | | CMF-RF | | | | | |
|---|---|---|---|---|---|---|---|---|---|---|---|---|---|---|---|---|---|---|---|---|---|---|---|---|---|
| Ring | Year | Angle | *r*-value | Oriented | Random | Inactive | *N* | Angle | *r*-value | Oriented | Random | Inactive | *N* | Angle | *r*-value | Oriented | Random | Inactive | *N* | Angle | *r*-value | Oriented | Random | Inactive | *N* |
| 17 white | 2022 | 85.06 | 0.217 | 7 | 0 | 3 | 10 | 349.38 | 0.203 | 9 | 0 | 2 | 11 | 104.81 | 0.751 | 6 | 0 | 0 | 6 | 335.06 | 0.313 | 8 | 0 | 0 | 8 |
| 19 white | 2022 | 345.46 | 0.245 | 14 | 2 | 1 | 17 | 164.76 | 0.093 | 12 | 4 | 0 | 16 | 153.73 | 0.121 | 11 | 4 | 0 | 15 | 251.00 | 0.222 | 5 | 4 | 0 | 9 |
| 22 violet | 2022 | 7.57 | 0.219 | 16 | 1 | 0 | 17 | 295.12 | 0.171 | 16 | 0 | 0 | 16 | 43.78 | 0.466 | 9 | 0 | 0 | 9 | 92.92 | 0.354 | 9 | 0 | 0 | 9 |
| 43 violet | 2022 | 205.84 | 0.229 | 9 | 1 | 0 | 10 | 87.35 | 0.152 | 13 | 1 | 0 | 14 | 95.63 | 0.163 | 6 | 0 | 0 | 6 | 29.26 | 0.819 | 5 | 4 | 0 | 9 |
| 44 white | 2022 | 82.82 | 0.279 | 10 | 0 | 6 | 16 | 261.95 | 0.185 | 9 | 0 | 4 | 13 | 152.69 | 0.385 | 6 | 1 | 5 | 12 | 51.21 | 0.721 | 7 | 0 | 2 | 9 |
| 58 red | 2022 | 170.34 | 0.167 | 18 | 7 | 0 | 25 | 242.45 | 0.206 | 8 | 5 | 1 | 14 | 79.53 | 0.752 | 5 | 4 | 0 | 9 | 25.16 | 0.32 | 5 | 4 | 0 | 9 |
| 61 orange | 2022 | 4.75 | 0.244 | 13 | 0 | 4 | 17 | 218.15 | 0.247 | 12 | 0 | 5 | 17 | 245.02 | 0.473 | 8 | 0 | 1 | 9 | 73.12 | 0.4 | 8 | 0 | 1 | 9 |
| 67 white | 2022 | 47.31 | 0.299 | 6 | 0 | 5 | 11 | 340.60 | 0.33 | 14 | 0 | 3 | 17 | 120.22 | 0.561 | 4 | 0 | 2 | 6 | 99.70 | 0.303 | 7 | 0 | 2 | 9 |
| 72 violet | 2022 | 25.57 | 0.318 | 6 | 0 | 5 | 11 | 322.26 | 0.323 | 7 | 0 | 7 | 14 | 342.95 | 0.311 | 8 | 0 | 4 | 12 | 1.03 | 0.68 | 5 | 1 | 3 | 9 |
| 75 violet | 2022 | 141.50 | 0.145 | 21 | 1 | 3 | 25 | 224.54 | 0.327 | 13 | 1 | 2 | 16 | 79.88 | 0.252 | 8 | 1 | 0 | 9 | 336.21 | 0.104 | 7 | 1 | 1 | 9 |
| 82 violet | 2022 | 162.07 | 0.351 | 10 | 2 | 2 | 14 | 302.87 | 0.455 | 4 | 0 | 3 | 7 | | | | | | | | | | | | |
| 88 violet | 2022 | 320.86 | 0.248 | 13 | 0 | 0 | 13 | 251.12 | 0.712 | 9 | 1 | 1 | 11 | 43.22 | 0.072 | 7 | 2 | 0 | 9 | 205.00 | 0.275 | 3 | 0 | 0 | 3 |
| 97 orange | 2022 | 73.22 | 0.665 | 12 | 1 | 0 | 13 | 317.89 | 0.056 | 14 | 0 | 0 | 14 | 45.28 | 0.294 | 5 | 4 | 0 | 9 | 69.08 | 0.647 | 5 | 4 | 0 | 9 |
| 99 orange | 2022 | 5.21 | 0.372 | 6 | 0 | 5 | 11 | 162.81 | 0.27 | 9 | 1 | 6 | 16 | 9.06 | 0.208 | 9 | 0 | 0 | 9 | 248.23 | 0.26 | 9 | 0 | 0 | 9 |
| 01 orange | 2023 | 334.69 | 0.136 | 15 | 0 | 0 | 15 | 89.52 | 0.09 | 9 | 0 | 0 | 9 | 73.60 | 0.384 | 5 | 1 | 0 | 6 | 200.76 | 0.309 | 8 | 1 | 0 | 9 |
| 08 orange | 2023 | 19.08 | 0.162 | 7 | 0 | 2 | 9 | 35.19 | 0.574 | 6 | 0 | 3 | 9 | 60.09 | 0.492 | 6 | 0 | 3 | 9 | 75.62 | 0.132 | 17 | 0 | 1 | 18 |
| 16 orange | 2023 | 280.75 | 0.042 | 5 | 0 | 1 | 6 | 337.27 | 0.573 | 6 | 0 | 3 | 9 | 114.33 | 0.095 | 4 | 0 | 2 | 6 | 58.93 | 0.861 | 4 | 0 | 2 | 6 |
| 23 orange | 2023 | 51.58 | 0.657 | 5 | 1 | 0 | 6 | 256.51 | 0.228 | 24 | 3 | 0 | 27 | 21.35 | 0.591 | 6 | 0 | 0 | 6 | 320.48 | 0.498 | 5 | 1 | 0 | 6 |
| 30 orange | 2023 | 55.17 | 0.464 | 7 | 0 | 2 | 9 | 224.46 | 0.471 | 7 | 2 | 0 | 9 | 360.00 | 0.386 | 5 | 1 | 0 | 6 | 263.97 | 0.297 | 5 | 4 | 0 | 9 |
| 32 orange | 2023 | 37.83 | 0.325 | 9 | 0 | 0 | 9 | 157.27 | 0.05 | 18 | 0 | 0 | 18 | 33.38 | 0.415 | 9 | 0 | 0 | 9 | 345.88 | 0.48 | 6 | 0 | 0 | 6 |
| 33 orange | 2023 | 4.38 | 0.23 | 15 | 0 | 0 | 15 | 303.94 | 0.487 | 9 | 0 | 0 | 9 | 112.58 | 0.295 | 8 | 0 | 1 | 9 | 359.39 | 0.492 | 12 | 0 | 0 | 12 |
| 41 violet | 2023 | 160.39 | 0.466 | 6 | 1 | 5 | 12 | 122.51 | 0.304 | 17 | 0 | 7 | 24 | 284.19 | 0.656 | 6 | 0 | 6 | 12 | 303.09 | 0.349 | 6 | 0 | 3 | 9 |
| 43 orange | 2023 | 320.75 | 0.184 | 4 | 0 | 5 | 9 | 134.70 | 0.104 | 8 | 0 | 7 | 15 | 24.51 | 0.603 | 6 | 0 | 3 | 9 | 61.62 | 0.678 | 13 | 0 | 4 | 17 |
| 45 white | 2023 | 8.89 | 0.527 | 4 | 0 | 2 | 6 | 206.17 | 0.291 | 8 | 0 | 1 | 9 | 27.69 | 0.539 | 8 | 0 | 1 | 9 | 310.00 | 0.333 | 3 | 0 | 0 | 3 |
| 48 orange | 2023 | 30.45 | 0.577 | 12 | 0 | 0 | 12 | 51.72 | 0.313 | 18 | 1 | 2 | 21 | 71.75 | 0.449 | 7 | 0 | 1 | 8 | 355.00 | 0.258 | 8 | 1 | 0 | 9 |
| 54 orange | 2023 | 125.21 | 0.473 | 10 | 0 | 2 | 12 | 332.46 | 0.248 | 8 | 0 | 1 | 9 | 132.91 | 0.264 | 8 | 0 | 1 | 9 | 345.74 | 0.511 | 7 | 0 | 2 | 9 |
| 55 orange | 2023 | 258.63 | 0.491 | 9 | 0 | 0 | 9 | 271.36 | 0.339 | 5 | 1 | 0 | 6 | 238.01 | 0.287 | 7 | 2 | 0 | 9 | 124.19 | 0.211 | 22 | 2 | 0 | 24 |
| 58 orange | 2023 | 40.27 | 0.3 | 6 | 3 | 0 | 9 | 56.45 | 0.175 | 10 | 2 | 3 | 15 | 85.53 | 0.502 | 5 | 0 | 1 | 6 | 195.30 | 0.26 | 12 | 3 | 6 | 21 |
| 97 white | 2023 | 50.31 | 0.496 | 10 | 2 | 0 | 12 | 339.61 | 0.599 | 5 | 1 | 0 | 6 | 52.98 | 0.525 | 6 | 3 | 0 | 9 | 356.39 | 0.486 | 7 | 2 | 0 | 9 |

### C Supplementary analysis

A non-significant orientation in the CMF-RF condition could either result from a) a lack of statistical power, or b) genuine disorientation caused by the presence of the RF field. Differentiating between these two is difficult, owing to the impossibility of proving a negative. In case a), a lack of statistical power would mislead one to conclude the birds' orientation behaviour was disrupted. Our obtained orientations should then show a degree of directedness that resembles oriented groups of birds from previous studies. In case b), the RF field exposure truly caused disorientation of the birds' behaviour and the degree of directedness should resemble disoriented groups of birds from previous studies (where disorientation was observed in both the NMF and CMF conditions). We looked into this by comparing the test conditions across the years separately, by bootstrapping the original data sets and using a randomization approach, where the bootstrap sampled from the combination of the present study and from previous studies of the CMF-RF condition in spring (0.1–100 kHz in [1] – no effect; 75–85 MHz in [2] – disorienting effect; 140–150 MHz in [3] – no effect). The methods and results of the bootstrap and randomization approach are described in the following, as well as the results of the statistical comparisons across years.

#### C.1 Extended comparison of experimental seasons and test conditions

As outlined in the main article, the NMF control conditions of both years were significantly oriented in the seasonally appropriate migratory direction (see SI-table 3). In the CMF controls, the groups of birds in both years only displayed a non-significant tendency towards W and NW, respectively (SI-table 3). The group orientation direction of the NMF control and CMF control were statistically different from each other in 2022 ($W_{(2)} = 11.25$, $p = 0.0036$), but not in 2023 (SI-table 4). While the group orientation turned with the 120°-counter-clockwise turn of the magnetic field in the CMF conditions (2022: 125°, 2023: 96°), this turn was only statistically significant for 2022 (SI-table 5).

Table 3: *Summary statistics of group orientation across the years (2022, 2023, and their combination), and across test conditions. Listed are the obtained sample size (N), the group circular mean ± angular standard deviation (s.d.), the group r-value, the groups' Z-score and p-value of the Rayleigh test, and the angular 95% confidence interval (CI) for significantly oriented subsets. The p-values of statistically significant Rayleigh tests are listed in bold.*

| Year | Condition | *N* | mean ± s.d. | *r*-value | *Z*-score | *p*-value | 95% CI |
|---|---|---|---|---|---|---|---|
| 2022 | NMF | 12 | 35° ± 57° | 0.5 | 3.01 | **0.0457** | 352° - 77° |
| | CMF | 9 | 270° ± 55° | 0.54 | 2.59 | 0.0718 | |
| | NMF-RF | 10 | 70° ± 58° | 0.49 | 2.41 | 0.0877 | |
| | CMF-RF | 12 | 42° ± 65° | 0.35 | 1.51 | 0.2257 | |
| 2023 | NMF | 11 | 45° ± 52° | 0.58 | 3.73 | **0.0202** | 8° - 82° |
| | CMF | 11 | 309° ± 66° | 0.34 | 1.3 | 0.2786 | |
| | NMF-RF | 14 | 51° ± 53° | 0.57 | 4.52 | **0.0085** | 17° - 85° |
| | CMF-RF | 14 | 340° ± 63° | 0.4 | 2.29 | 0.1001 | |
| Combined | NMF | 23 | 40° ± 55° | 0.54 | 6.65 | **0.0009** | 12° - 68° |
| | CMF | 20 | 287° ± 62° | 0.41 | 3.3 | **0.0348** | 245° - 329° |
| | NMF-RF | 24 | 58° ± 56° | 0.53 | 6.72 | **0.0008** | 30° - 87° |
| | CMF-RF | 26 | 6° ± 66° | 0.33 | 2.78 | 0.0606 | |

In the presence of 110-120 MHz broadband RF fields of intensity $B_{rms}$ = 2.1 nT, the 2022 cohort of birds displayed non-significant tendencies towards NE both in the NMF-RF and CMF-RF condition (SI-table 3). The orientations of the 2022 cohort of birds in the NMF-RF and CMF-RF was not statistically different from each other ($W_{(2)} = 1.29$, $p = 0.5257$) and they showed no statistically significant (angular difference: 28°, $V = -0.01$, $\mu = -0.07$, $p = 0.526$) change in the group mean orientation. The lack of significant group orientations and no clear rotation of the orientation in response to the 120°-CCW turned magnetic field could indicate a disorienting effect of the used broadband RF fields.

Table 4: *Summary statistics of Mardia-Watson-Wheeler (MWW) across test conditions and years. Listed are the W-scores with 2 degrees of freedom (d.f.) and p-values. The p-values of statistically significant MWW Tests are listed in bold.*

| Year | Comparison | $W_{(d.f.\,=\,2)}$ | *p* |
|---|---|---|---|
| 2022 | NMF vs. CMF | 11.25 | **0.0036** |
| | NMF-RF vs. CMF-RF | 1.29 | 0.5257 |
| | NMF vs. NMF-RF | 1.68 | 0.432 |
| | CMF vs. CMF-RF | 5.58 | **0.0615** |
| 2023 | NMF vs. CMF | 4.01 | 0.1345 |
| | NMF-RF vs. CMF-RF | 6.31 | **0.0426** |
| | NMF vs. NMF-RF | 1.97 | 0.3728 |
| | CMF vs. CMF-RF | 0.93 | 0.6288 |
| Combined | NMF vs. CMF | 10.75 | **0.0046** |
| | NMF-RF vs. CMF-RF | 4.93 | 0.0848 |
| | NMF vs. NMF-RF | 4.25 | 0.1192 |
| | CMF vs. CMF-RF | 5.25 | 0.0725 |

Table 5: *Summary statistics of V-Test results across test conditions and years. Listed are the V- and μ-scores, and p-values for the V-Test of the mean orientation data in the CMF condition against the group mean data of the NMF condition (adjusted for the 120°-counter-clockwise rotation of the magnetic field). The p-values of statistically significant V-Tests are listed in bold.*

| Year | Comparison | *V* | *μ* | *p* |
|---|---|---|---|---|
| 2022 | NMF vs. CMF | 0.53 | 2.27 | **0.0105** |
| | NMF-RF vs. CMF-RF | -0.01 | -0.07 | 0.526 |
| 2023 | NMF vs. CMF | 0.31 | 1.47 | 0.0712 |
| | NMF-RF vs. CMF-RF | 0.27 | 1.41 | 0.0801 |
| Combined | NMF vs. CMF | 0.4 | 2.55 | **0.005** |
| | NMF-RF vs. CMF-RF | 0.12 | 0.89 | 0.1878 |

According to our bootstrap results testing for significant differences in directedness (r-values), the group directedness of the control NMF group of 2022 fell within the bootstrapped *r*-value CI of the NMF-RF, and vice versa ($r_{\text{NMF-RF}}$ = 0.491, $r$-$\text{CI}_{\text{NMF-RF}}$ = 0.188 - 0.811; $r_{\text{NMF}}$ = 0.501, $r$-$\text{CI}_{\text{NMF}}$ = 0.174 - 0.856; SI-table 7A and B). Furthermore, at least 37.5% of the 100,000 bootstrap samples were as directed and oriented in a similar direction as the respective other condition. Thereby, the NMF and NMF-RF of 2022 were not more, or less, significantly directed than the other.

In the NMF-RF of 2023, with RF fields of the intensity of $B_{\text{rms}}$ = 1.8 nT, the birds oriented significantly in the seasonally appropriate NE migratory direction (SI-table 3). The group orientation in the CMF-RF of 2023 tended roughly towards magnetic East, without statistical significance (SI-table 3). While the NMF-RF and CMF-RF of 2023 were statistically different from each other ($W_{(2)} = 6.31$, $p = 0.0426$), their difference of 71° in the group mean direction turned out not to be statistically significant ($V = 0.27$, $\mu = 1.41$, $p = 0.0801$).

The group orientations in the NMF-RF of both years were directed towards NE, similar to their respective controls, and did not differ from their respective controls (see SI-table 4). As stated in the main article, there was no statistical difference between the NMF-RF conditions across the years ($W_{(2)} = 0.58$, $p = 0.7487$). Similarly, the CMF-RF conditions did not differ statistically from their controls (see table 4), and no difference was found in the comparison of the years ($W_{(2)} = 1.48$, $p = 0.4769$).

The variances of *r*-values between the test conditions did not differ significantly across the years (Levene's Test: $F_{(7,85)} = 1.01$, $p = 0.4305$). The consistency of the birds' directedness therefore was not affected by the migratory season or condition (magnetic field or presence of broadband RF field). We also found no differences between the years and test conditions in the variance of the birds' orientation angles (SI-table 6). Therefore, neither year, nor test condition had a statistically significant effect on how the birds' orientations spread around the group mean. However, we found that the individual orientations were significantly more spread (showed larger angular variation) around the group mean in the CMF-RF condition compared to both NMF conditions (Levene's Test and post-hoc Tukey HSD; SI-table 6 and SI-figure 2).

Table 6: *Result statistics of Levene's test of variance and over the test condition ('Cond') by year ('Year') and combined. Listed are the F-statistic and degrees of freedom (F(d.f.)), sum of squares (SS), mean of squares (MS). Below: p-values of the post-hoc Tukey HSD result for the combined years. The p-values of statistically significant terms or tests are listed in bold.*

| Levene's Test | Term | SS | MS | *F* (*d.f.*) | *p* |
|---|---|---|---|---|---|
| | Cond | 557.9 | 186 | 0.101 (3) | 0.959 |
| | Year | 3735.1 | 3735.1 | 2.037 (1) | 0.1572 |
| | Cond:Year | 5014.9 | 1671.6 | 0.912 (3) | 0.439 |
| | Residuals | 155882.5 | 1833.9 | — (85) | |
| Combined | Cond | 25163.7 | 8387.9 | 3.863 (3) | **0.012** |
| | Residuals | 193267.1 | 2171.5 | — (89) | |

| Tukey HSD | CMF | CMF-RF | NMF |
|---|---|---|---|
| CMF-RF | (n.s.) 0.1825 | | |
| NMF | (n.s.) 0.9253 | * **0.0331** | |
| NMF-RF | (n.s.) 0.8386 | * **0.0166** | (n.s.) 0.9965 |

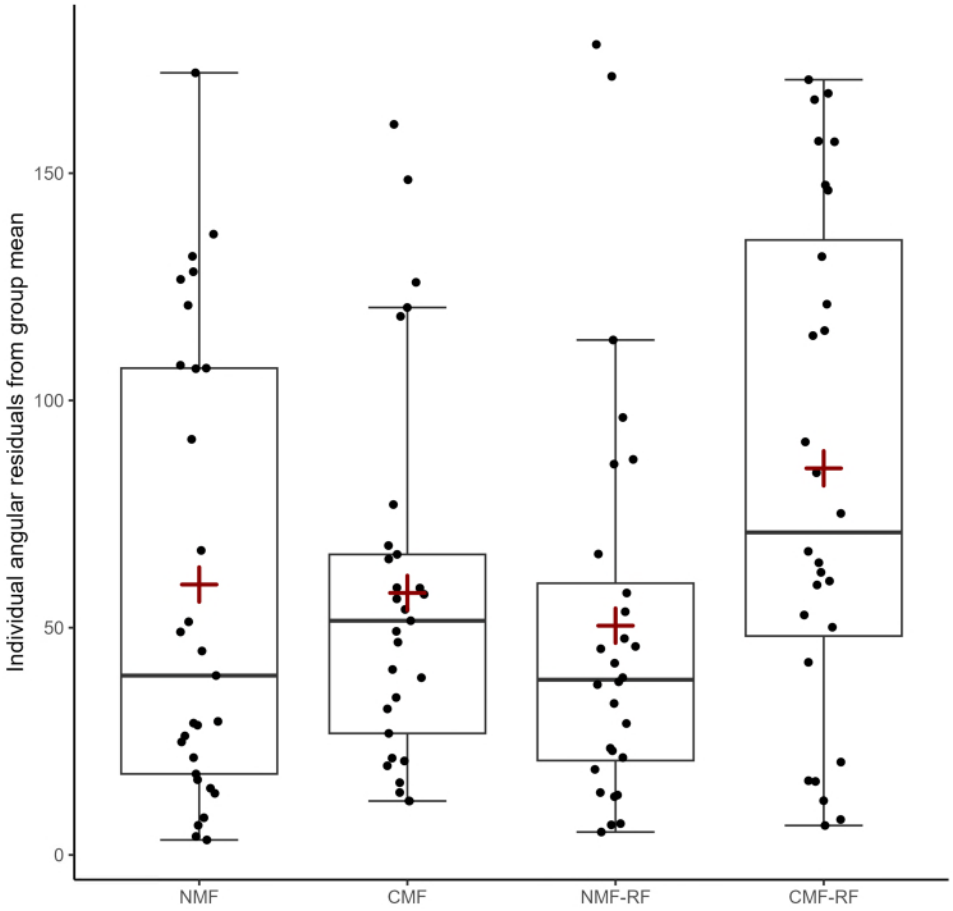


Figure 2: *Boxplots of the absolute residuals from the condition-specific group orientation. Each data point represents the absolute difference between an individual's orientation and the group mean of a given condition in degrees. The boxes range from Q1 (the first quartile) to Q3 (the third quartile) of the data distributions, and the range represents the IQR (interquartile range). Medians are indicated by lines across the boxes. The means are indicated by the red cross. The whiskers extend between the most extreme data points, excluding outliers, which are defined as being outside 1.5 IQR above the upper quartile and below the lower quartile.*

#### C.2 Resampling analysis (Bootstrap and Randomization) of behavioural data

Bootstrapping and randomization are both methods utilizing re-sampling. The process common to both methods will be described here, while the method-specific differences are described in the respective method section in the following.

For both methods, *n* samples (where *n* is the number of birds providing a mean orientation value in the given condition) are randomly picked from a given data set with replacement (meaning the same value could be sampled multiple times). In the present analysis, the individual orientation results for every bird tested in the given test condition is sampled. From these *n* samples a parameter of choice is calculated. The parameter of choice for both methods is the group mean vector length (the *r*-value, or so-called directedness), which represents how consistently the group of birds oriented in the same direction in the given condition. The process of random sampling with replacement is then repeated 100,000 times (a common number for re-sampling methods). From the 100,000 values for *r*, the 95% confidence intervals (CI) was estimated by ordering the 100,000 *r*-values from the lowest to the highest. The lower bound for the 95% CI for *r* is the value at position 2500 and the upper bound is the value at position 97500. Depending on how much the 95% CI of two re-sampled distributions overlap, or if the experimentally observed value lies within the 95% CI, similarities or differences in the sampled data sets can be inferred.

##### C.2.1 Bootstrap – Method

To investigate the potential disorienting effect of our experimental broadband 110–120 MHz RF-field, we used a common bootstrap approach [2–6]. With the bootstrapping, we wanted to test whether the directions in the broadband RF condition were significantly more spread out (did they show a significantly lower *r*-value) than the respective control condition. In the following, we describe the process for the comparison of the CMF-RF and the CMF condition combined across years. First, we merged the data from both years for the CMF-RF and CMF condition. The previously described re-sampling process was then applied to the combined CMF-RF data with *n* being the sample size of the CMF condition. In addition to the 95% CI of the *r*-values, the 95% CI of the group mean angles were calculated. For this purpose, the group mean angle of each re-sampling repetition was calculated. The

100,000 group mean angles were ordered in an ascending manner starting at the observed group mean angle minus 180° and ending at the observed group mean angle plus 180°. To consider the circular nature of the data, 360° was added to any angle smaller than the observed group mean angle minus 180°. If the parameter of interest, e.g. the *r*-value of the CMF condition ($r_{CMF}$-value), lies outside of the 95% CI of the bootstrapped data, this indicates that the underlying distributions of CMF and CMF-RF condition are unlikely to originate from the same base distribution. A *p*-value can be calculated by dividing the number of bootstrapped *r*-values exceeding the $r_{CMF}$-value by the total number of sampling repetitions (100,000). We thereby acquired the probability of how likely it is that the data of the CMF-RF condition (bootstrapped CMF-RF *r*-values) are as directed as the CMF condition (> $r_{CMF}$-value). We then continued, by calculating how many of the sampling repetitions are at least as directed as the observed CMF data *and* also oriented in a direction lying within the 95% CI for the mean direction of the CMF condition. Therefore, the sampling repetitions exceeding the $r_{CMF}$-value were further filter for whether the fall between the angular 95% CI of the CMF condition. Dividing this number by the number of sampling repetitions (100,000) gives us the *p*-value for whether the bootstrapped CMF-RF data are as directed and oriented as the CMF data.

**C.2.2 Bootstrap – Results**

The observed group mean directedness (*r*-value) of both CMF conditions fell well between the bootstrapped 95% CI of the respective mean *r*-values (compare the observed *r*-values in SI-table 7A with the *r*-value CIs in SI-table 7B). A large majority of the bootstrap iterations (CMF: 83.6% of 100,000 bootstrap iterations; CMF-RF: 38.7%) were at least as directed as the corresponding CMF condition with or without RF fields. According to our bootstrap analysis, both CMF conditions cannot be considered significantly different in directedness (see *p*-values for "> compared *r*-value" in SI-table 7C). This is visualized in the large portions of the heat maps exceeding the compared *r*-value (dashed lines in SI-figure 3). We further tested how many of the bootstrap iterations of the CMF-RF data were as directed (same or larger *r*-value) as the CMF data and fell within the angular CI of the CMF condition. The obtained values confirm the insufficient turn in the CMF-RF condition since only a few bootstrap iterations of the CMF-RF data were as directed and fell within the angular CI of the respectively other

CMF group (CMF-RF: 1.7% ; CMF: 3.5%). These low percentages translated to significant differences between the CMF and CMF-RF condition (see *p*-values for "> compared *r*-value & in angular CI" in SI-table 7B). The visualization of the bootstraps in SI-figure 3 illustrate how both conditions are similarly directed, but with different orientation.

Table 7: *Bootstrap results for the comparison of the CMF conditions absence and presence of a 110–120 MHz broadband RF field. The bootstrapped condition was resampled (N = 100,000) with replacement with the sample size of the compared condition. Section (A) lists the parameters of the compared condition ('Parameters from'), which were used for the bootstrap analysis of the bootstrapped condition ('Data from'). The given parameters are sample size (N), r-value, and angular confidence interval (CI), which resulted in the bootstrapped CIs for group mean angle and r-value obtained for each test condition. Section (B) lists the results of the comparison of the bootstraps with the observed data: the number and percentage of bootstrap iterations (N), and corresponding p-values exceeding the compared r-value; the same statistics for iterations that exceed the compared r-value and fell in-between the compared angular CI bounds. The p-values of statistically significant comparisons are listed in bold.*

| (A) | | Parameters | | | | Bootstrapped CIs | |
|---|---|---|---|---|---|---|---|
| Year | Parameters from | angular CI | *r*-value | *N* | Data from | group mean direction | *r*-value |
| 2022 | NMF-RF | 22.3° - 118.2° | 0.491 | 10 | NMF | 353° - 97° | 0.174 - 0.856 |
| | NMF | 351.8° - 77.3° | 0.501 | 12 | NMF-RF | 22.5° - 116.6° | 0.188 - 0.811 |
| Combined | CMF-RF | 320° - 52.6° | 0.327 | 26 | CMF | 247.3° - 326.1° | 0.213 - 0.63 |
| | CMF | 245.4° - 329° | 0.406 | 20 | CMF-RF | 299.2° - 63.7° | 0.105 - 0.627 |

| (B) | | > compared *r*-value | | > compared *r*-value & in angular CI | |
|---|---|---|---|---|---|
| Year | Comparison | *N* | *p* | *N* | *p* |
| 2022 | NMF vs. NMF-RF | 62327 (62.3%) | 0.6233 | 43227 (43.2%) | 0.4323 |
| | NMF-RF vs. NMF | 58534 (58.5%) | 0.5853 | 37494 (37.5%) | 0.3749 |
| Combined | CMF vs. CMF-RF | 83552 (83.6%) | 0.8355 | 3539 (3.5%) | **0.0354** |
| | CMF-RF vs. CMF | 38724 (38.7%) | 0.3872 | 1688 (1.7%) | **0.0169** |

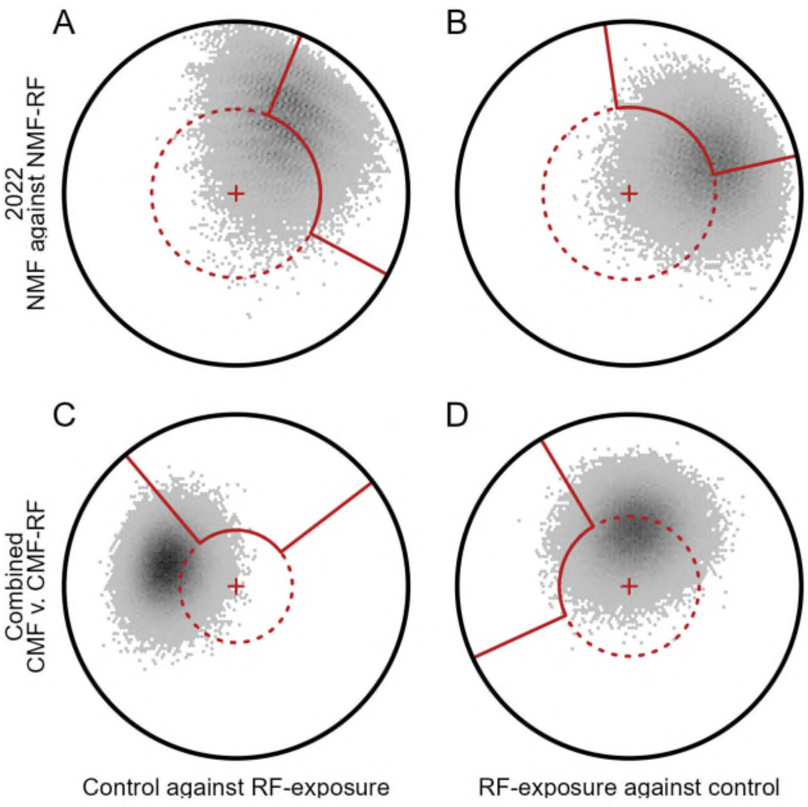


Figure 3: *Circular heatmap representation of the mean direction and directedness of each bootstrap sampling (N = 100,000). The data of the 2022 NMF (A, B) and combined CMF condition (C, D) in the absence (A, C) and presence of a 110–120 MHz broadband RF field (B, D), were bootstrapped with the sample size of the other condition (e.g., the data of the control is bootstrapped with the sample size of the RF condition). The bootstrapped data are plotted with the 95% confidence intervals (solid lines) and group directedness (r-values; dashed circles) of the other condition (e.g., the dashed circle in A represents the r-value of the RF condition in B). The darker an area, the more bootstrap iterations resulted in an angle and directedness in that area. Number of heatmap bins = 100.*

Besides comparing the non-significant orientation of the combined CMF-RF to its CMF control, we applied the same approach to the non-significant NMF-RF and NMF control of 2022. The observed group *r*-value of either condition fell well within the *r*-value CI obtained by bootstrapping (SI-table 7A and B). A large majority of the bootstrap samples for the NMF and NMR-RF condition in 2022 were at least as directed as the respective control or broadband RF condition (SI-table 7C). The bootstrap comparison indicates that the NMF and NMF-RF condition of 2022 are similarly directed in a similar direction, and not significantly different from each other.

##### C.2.1 Randomization – Method

To further explore the effect of our broadband RF fields on the birds' orientation, we also compared our CMF-RF data (separated by year and with combined years) to previous CMF conditions under broadband RF fields conducted in spring seasons [1–3]. To do so, we used a randomization approach, which differs from bootstrapping in the composition of data from which is sampled. For the randomization approach, the random sampling was applied to a combination of our CMF-RF data and the RF data of a previous study [1–3]. As the analysis is focused on the similarity in group directedness *r* between the studies, the (minor) differences in orientation between the studies had to be transformed in one common direction. For transformation, the group mean direction of each data set was subtracted from the individual data points. Thereby, the distribution of the data points was conserved and all group mean directions were centred on 0°. The re-sampling process was then applied to each transformed combination of data sets, with 26 sampled data points per repetition (the sample size of the present CMF-RF condition).

Thereby, we sought to see whether the *r*-value of the present CMF-RF condition under 110–120 MHz broadband RF fields resembled previously tested broadband conditions leading to disorientation (75–85 MHz: [2]) or to non-disorientation of the birds' magnetic compass orientation (0.1–100 kHz: [1]; 140–150 MHz: [3]). Similar to bootstrapping, the 95% CI and median for the group *r*-value were calculated for each sampled combination. For illustrative purposes we plotted the distributions of the bootstrapped *r*-values for the CMF-RF condition of the present and the previous studies in SI-figure 4.

##### C.2.4 Randomization – Results

From the bootstrapped *r*-values of the original data from previous studies, the 95% confidence intervals (CI) and median can be separated into two groups (red lines in SI-figure 4). Broadband RF fields with no effect (SI-figure 4; 0.1–100 kHz [1] and 140–150 MHz [3]) produced higher *r*-value distributions with overlapping CIs (see "Bootstrapping" SI-table 8). By contrast, broadband RF fields with a disorienting effect (SI-figure 4; 75–85 MHz [2]) produced *r*-value distributions closer to zero. Notably, the CIs of the previous studies with and without a disorienting effect on the birds orientation do not overlap, implying that the distributions distinctly represent disoriented [2] and oriented data [1,3]. The distribution and CI of the bootstrapped *r*-values of the present study (regardless of the separation or combination by year) overlap with all the distributions and CIs of the previous studies (blue lines in SI-figure 4 and SI-table 8A). In other words, they fall in-between the previous oriented and disoriented distributions. This is an indication that the 110-120 MHz fields might have been distractive, but not destructive, i.e. they seem to be semi-oriented.

Table 8: *Results obtained by bootstrapping and randomization of previous and present orientation data in the CMF-RF condition under broadband RF fields. Section (A): The frequency range of the broadband fields are separated by study and year (left-most column) and the bootstrapped r-value median and 95% confidence intervals (CI) for the group r-value are given. The results are visualized in SI-figure 4 'Original data'. Section (B): The CMF-RF condition of the present study (separated by year) and the observed r-value are listed next to the CMF-RF condition of the previous studies with which the data was combined before randomization. The r-value median and 95% CI resulting from randomization are given for each combination. The results are visualized in SI-figure 4 'Randomized'. Either, the bootstrapped condition and the randomization combination were resampled (N = 100,000) with replacement with a sample size of n = 26.*

(A) Bootstrapping

| Frequency broadband | Bootstrapped *r*-value | |
|---|---|---|
| | Median | 95% Confidence interval (CI) |
| 0.1–100 kHz | 0.603 | 0.394 - 0.783 |
| 75–85 MHz | 0.176 | 0.36 (one-sided) |
| 140–150 MHz | 0.627 | 0.47 - 0.76 |
| 110–120 MHz (2022) | 0.376 | 0.107 - 0.639 |
| (2023) | 0.425 | 0.164 - 0.669 |
| (Combined) | 0.357 | 0.117 - 0.591 |

(B) Randomization

| Present data from year | present *r*-value | Randomized with data from | Randomization *r*-value | |
|---|---|---|---|---|
| | | | Median | 95% CI |
| 110–120 MHz (2022) | 0.354 | 0.1–100 kHz | 0.502 | 0.245 - 0.726 |
| | | 75–85 MHz | 0.215 | 0.043 - 0.456 |
| | | 140–150 MHz | 0.502 | 0.255 - 0.711 |
| 110–120 MHz (2023) | 0.404 | 0.1–100 kHz | 0.517 | 0.275 - 0.731 |
| | | 75–85 MHz | 0.232 | 0.048 - 0.474 |
| | | 140–150 MHz | 0.518 | 0.286 - 0.719 |
| 110–120 MHz (Combined) | 0.327 | 0.1–100 kHz | 0.445 | 0.202 - 0.664 |
| | | 75–85 MHz | 0.244 | 0.052 - 0.481 |
| | | 140–150 MHz | 0.44 | 0.206 - 0.652 |

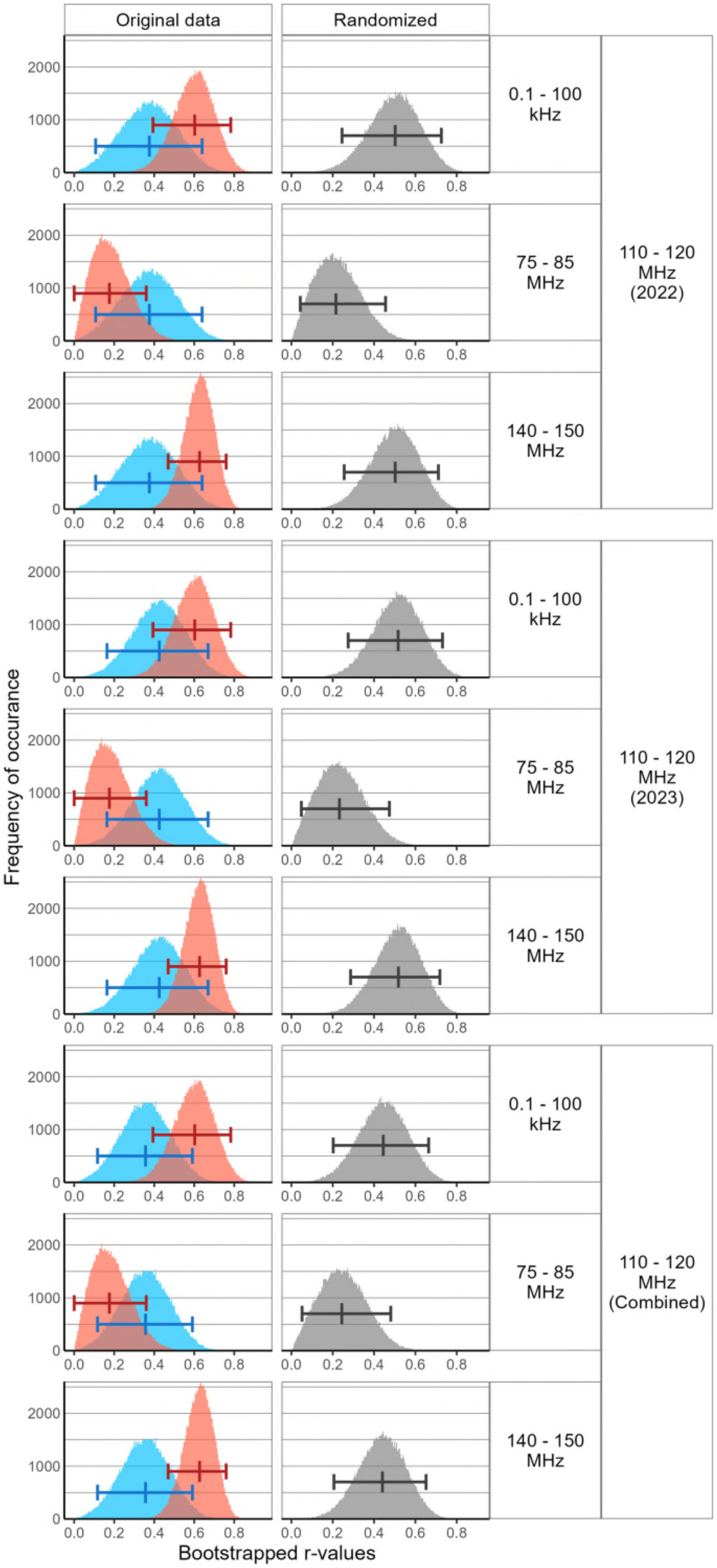
Original data
Randomized
0.1 - 100 kHz
75 - 85 MHz
140 - 150 MHz
110 - 120 MHz (2022)
110 - 120 MHz (2023)
110 - 120 MHz (Combined)
Frequency of occurance
Bootstrapped r-values
0
1000
2000
0.0
0.2
0.4
0.6
0.8

Figure 4: *Expanded version of figure 3 in the main article. Histograms of the bootstrapped r-values of the original data of the CMF condition in the presence of broadband RF fields in the present study in the frequency range 110–120 MHz (blue) side-by-side with the previous CMF conditions in other broadband RF fields (red), and the randomized bootstrapped r-values of combinations of the data of our present and the respective previous CMF condition with broadband RF fields present (grey). Rows: non-disruptive broadbands from 0.1–100 kHz in Kobylkov et al. 2019; disruptive broadbands from 75–85 MHz in Leberecht et al. 2022; non-disruptive broadbands from 140–150 MHz in Leberecht et al. 2023. The columns show the bootstrapped 'Original Data' (red and blue histograms) and bootstrapped 'Randomization' (grey histograms) r-values separated by year (2022, 2023, and their combination). Each histogram represents 100,000 bootstrap iterations, each with 26 samples (sample size for our combined 110–120 MHz CMF-RF condition) randomly picked with replacement, from which the r-value was computed. The whiskers indicate the median and bootstrapped 95% confidence intervals. Number of bins = 200.*

After randomized re-sampling of each of the previous studies data combined with our data, the shape of the distribution clearly changed (compare red and blue histograms in SI-figure 4 to the grey histograms), resembling a mixture of the bootstrapped *r*-values of the original data sets. In all combinations, the CIs overlap each other (see SI-table 8B). The CIs of all randomized distributions overlapped (SI-table 8B and grey lines in SI-figure 4), unlike their original CIs (SI-table 8A and red lines in SI-figure 4). We therefore cannot claim that the distribution of *r*-values found under broadband 110–120 MHz RF is significantly different from oriented or disoriented the previous data sets.

#### C.3 Summary

To summarize, from our bootstrap analysis of the *r*-values, we can conclude that the birds as a group orient equally well in the condition-specific group direction, regardless whether RF fields are present or not. Thus, the randomization analysis suggest that the orientation distributions found under broadband 110-120 MHz RF seems to fall in-between random and oriented samples, as it is not significantly different from either the previously reported oriented groups or the previously reported disoriented groups in Leberecht et al. [2,3] or Kobylkov et al. [1].
